# PackDock: a Diffusion Based Side Chain Packing Model for Flexible Protein-Ligand Docking

**DOI:** 10.1101/2024.01.31.578200

**Authors:** Runze Zhang, Xinyu Jiang, Duanhua Cao, Jie Yu, Mingan Chen, Zhehuan Fan, Xiangtai Kong, Jiacheng Xiong, Zimei Zhang, Wei Zhang, Shengkun Ni, Yitian Wang, Shenghua Gao, Mingyue Zheng

## Abstract

Structure-based drug design (SBDD) relies on accurate knowledge of protein structure and ligand-binding conformations. However, most of the static conformations obtained by advanced methods such as structural biology and de novo protein folding algorithms often don’t meet the needs for drug design. We introduce PackDock, a flexible docking method that combines “conformation selection” and “induced fit” mechanisms in a two-stage docking pipeline. The core module of this method is PackPocket, which uses a diffusion model to explore the side-chain conformation space in ligand binding pockets, both with or without a ligand. We evaluate our method using several tests that reflect real-world application scenarios. (1) Side-chain packing and Re-docking experiments validate the ability of PackDock to predict accurate side-chain conformations and ligand conformations. (2) Cross-docking experiments with *apo* and non-homologous ligand-induced *holo* structures align with real docking scenarios, demonstrating PackDock’s practical value. (3) Docking experiments with hypothetical models show that PackPocket can potentially conduct SBDD starting from protein sequence information only. Additionally, we found that PackDock can identify key amino acid conformation changes, which may provide insights for lead compound optimization. We demonstrate PackDock can accurately predict the complex conformations in various application scenarios, by combining the conformation selection theory and the induced fit theory, and by using the ability of PackPocket to accurately predict the side chain conformations in the pocket region. We believe this method can improve the usability of existing structures, providing a new perspective for the SBDD community.

## Introduction

Over the past few decades, Structure-Based Drug Design (SBDD) has been a primary approach for drug design with identified molecular targets ^[1-3]^. Owing to the rapid advancement of structural biology and de novo protein folding algorithms, three-dimensional protein structures are now more accessible^[4-11]^. However, the existing protein structures often don’t meet the needs of SBDD. For example, *apo* structures and *holo* structures with non-homologous ligands may lead to incorrect judgments of ligand binding modes, and not all predicted structures by protein folding are suitable for SBDD^[12, 13]^. This is primarily because the existing structures are static and do not account for the complex structural flexibility that proteins, which adapt their internal movements to their respective molecular binders^[13-15]^. Therefore, there is a high demand for methods that take into account the structural flexibility of complexes, which would overcome these limitations and broaden the scope of applicable targets for SBDD.

The dynamic protein-ligand binding process is commonly described by two classical mechanisms: (1) Conformational selection: a ligand selectively binds to only a few existing receptor conformations. This binding disrupts the dynamic equilibrium of receptor conformations, causing more conformations to collapse towards the selected one ^[16-18]^. (2) Induced Fit: a ligand binds to the main free conformation of the receptor, which then undergoes a conformational change to produce a more tightly bound complex conformation.^[19-21]^. Protein conformational changes during binding processes in different systems may follow different mechanisms or have both mechanisms simultaneously^[22-25]^. When using protein structures other than *holo*, take no account of protein flexibility may result in failure of complex structure prediction^[15, 26]^ or unsatisfactory performance of high-throughput virtual screening for new ligands^[27, 28]^. Traditional methods rely on molecular dynamics (MD) to consider protein flexibility, while they are time-consuming and not suitable for large-scale ligand virtual screening scenarios. Therefore, we need an efficient method that can account for both the conformational selection and induced fit theories of protein conformational changes.

In most cases of rigid docking failures, adjusting the side chain direction of the *apo* protein structure enables successful docking of ligands. These ligands are usually positioned close to their crystal structure conformation, typically within 2.5 Å RMSD(Root Mean Square Deviation) when compare to their crystal conformation^[29, 30]^. Numerous experiments have shown that the main chain movement from the *apo* to the *holo* structure is generally minimal (typically less than 1 Å). Most docking failures are due to changes in the side chain conformation around the pocket ^[31-33]^. Therefore, a strategy that balances accuracy and complexity might involve transitioning from full-flexibility docking to accommodating only local flexibility. This is often viewed as an effective approach for flexible docking, and most of the existing flexible docking algorithms utilize rotamer library databases and heuristic searches following this strategy. However, the computational time increases exponentially with the number of flexible side chains. Moreover, manual specification of critical side chains is required, limiting its application in SBVS. Examples include methods like *Vina*_*flex*_ ^[34]^, *AutoDockFR* ^[35]^ (*AutoDock* for Flexible Receptors). In addition, IFD-MD^[36]^ constructs a flexible docking pipeline that can slightly adjust the main chain of the pocket region through empirical scoring functions, implicit solvent force field energies etc., but requires more computational resources. Among deep learning-based methods, some only consider the coarse-grained *C*_*α*_ of the protein, implicitly considering some protein flexibility when obtaining the ligand pose, but they fail to obtain the atomic details of the protein-ligand complex structure required for SBDD. For instance, EDM-dock^[37]^ rebuilds complex conformation through protein-ligand distance maps. Equibind^[38]^, E3bind^[39]^, Uni-Mol^[40]^ predicts ligand three-dimensional coordinates through equivariant neural networks. Additionally, DiffDock^[41]^ simplifies the docking problem to the generation of ligand rotation, translation, and torsion angles. Recently, FlexPose^[42]^ expanded the prediction ability of flexible docking methods on protein side chain conformation by directly predicting the three-dimensional coordinates of the complex. However, current deep learning methods all have their own limitations. For example, they can generate unreasonable ligand conformations due to the insufficient learning of physical constraints, or spatial conflicts can occur between the ligand and the protein due to not considering the side chain conformation. This makes it impossible to provide reasonable protein-ligand interaction information, resulting in inapplicability to SBDD. Therefore, we urgently need a flexible docking method that can avoid these limitations, efficiently consider the conformational changes of protein side chains, and directly serve SBDD.

In this study, we developed a flexible docking pipeline, PackDock, which combines the benefits of conventional and DL-based methods. It takes into account both the conformation selection hypothesis and the induced fit hypothesis in receptor-ligand binding. Our method’s core module is PackPocket, a side chain packing model based on the diffusion model. This model predicts the various conformational changes of protein side chains in “free” and ligand “bound” states, and streamlined the problem of side-chain flexibility modeling by predicting the side chain torsional angles. We then conducted extensive tests on our method across multiple SBDD tasks. Firstly, to evaluate the feasibility of our method, we tested it with side-chain recovery and flexible re-docking experiments. In the side-chain recovery experiments, PackPocket outperforms all existing methods in the predicting accuracy of side-chain in pocket area under different scenarios. In the flexible re-docking experiments, PackDock performs significantly better than the corresponding flexible docking methods and is comparable to the corresponding rigid docking methods using *holo* structures. Then, to evaluate the practicality of our method, we tested it with CrossDock test by using *apo* and *holo* for flexible docking tests, aligning the test with real docking scenarios. Our results show that PackDock significantly outperforms existing docking methods. Next, to test the potential of our method for drug design from sequences, we used the docking test set with non-experimental structures (predicted by AF2). PackPocket significantly improved the performance of the AF2 predicted structures in related tasks. Finally, by visualizing the side chain conformations generated during the docking process, we found that PackDock can capture the potential side chain conformational changes of key amino acids. This proves the validity of its generated side chain conformations and may provide new insights for compound optimization. In conclusion, we demonstrated the practical utility of PackDock through a series of validations that align well with real-world applications. We hope this study offers a practical strategy for the community to push beyond the limitations of existing structures for SBDD applications.

## Results

### Overview of PackDock

When knowledge about protein-ligand binding mode is unavailable, flexible docking can be more beneficial for drug design. However, conventional flexible docking algorithms rely on inefficient heuristic searches against rotamer library databases, and current deep learning methods could lead to unreasonable complex conformations due to a lack of physical constraints^[43]^. To address these issues, we propose a novel docking pipeline, PackDock, which combines the advantages of deep learning and conventional rigid docking algorithms and eliminates unnecessary iterative processes. The aim is to improve the efficiency of flexible docking without compromising atomic details of the protein-ligand complex structure needed for SBDD.

PackDock, as shown in **Fig. 1a**, integrates sampling across two stages. It considers both the conformational selection and induced fit mechanisms simultaneously. This design eliminates the need for extensive computational iterations typically involved in multi-step search processes. During the conformational sampling phase of the protein pocket, we leverage the conditional generation advantage of diffusion models to build two separate side chain packing models. These models allow for the simultaneous consideration of potential pocket conformations, both with and without the ligand present in the pocket. During the conformational sampling phase of the ligand, we utilized a commonly employed conventional docking tool, *Vina* ^[34]^. Notably, PackDock can accommodate any docking algorithms, has great adaptability, and may benefit from using other docking algorithms to enhance flexible docking performance.

**Fig. 1.**
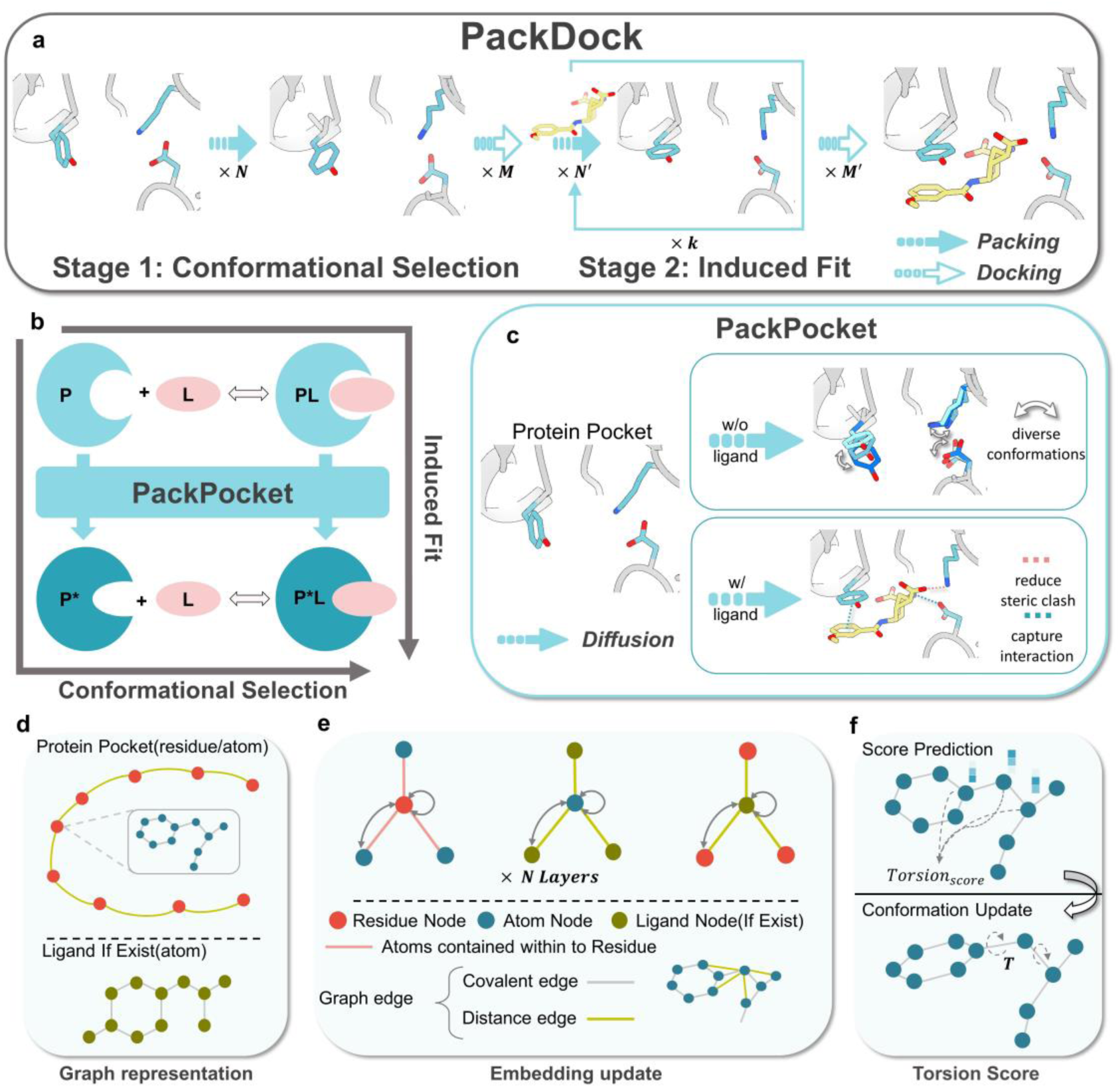
Overview of PackDock. **(a)** PackDock operates in two stages. Stage 1 uses a diffusion model PackPocket to explore empty pocket side-chain conformations, following the conformational selection hypothesis. In Stage 2, the diverse conformations obtained are used as receptor for docking. The conformation of the docked ligand is utilized to model the influence of the ligand in pocket conformation during binding, following the induced fit hypothesis. This process results in dynamic receptor-ligand complex conformations. **(b)** PackPocket can consider both the conformational selection mechanism and the induced fit mechanism simultaneously. **(c)** Diffusion process of the PackPocket module. This can be done with or without a given ligand. **(d)** Atom Graph and Ligand Graph of the Pocket Region. The atom graph of the pocket region comprises residue nodes, each containing atomic nodes for that residue. If a ligand is present, its atoms are used as nodes. **(e)** Embedding Update. The information of residue nodes is updated by atomic nodes of the corresponding residue. The information from ligand atom nodes is used as input to update the corresponding residue nodes and atomic nodes. Orange edges represent atoms containing within residue, grey edges represent covalent edges, and green edges represent distance edges. **(f)** Torsion Score. The score model updates the torsional conformation by atomic level embedding.

Stage 1: The first step is to simulate the conformational selection process. **Fig. 1a** shows that, in the absence of a ligand, the first pocket side-chain packing model can generate ***N*** conformations of the pocket in its “free” state by using the diversity-generating ability of the diffusion model. Each pocket conformation can sample ***M*** possible protein-ligand binding modes by docking. Finally, ***M*N*** possible complex conformations can be obtained to simulate the conformation selection process.

Stage 2: The second step is to simulate the induced-fit process. **Fig. 1a** shows that, in the condition of using the multiple ligand conformations obtained in the first stage, for each ligand conformation, the second pocket side-chain packing model can generate ***N***^′^ conformations of the pocket in the ligand “bound” state by using the conditional generation ability of the diffusion model. Each pocket conformation can also sample ***M***^′^ possible protein-ligand binding modes by docking to complete one iteration. The above process can be repeated ***k*** times, finally we can obtain ***k***(***NM***)(***N***^′^***M***^′^) different complex conformations. In this work, we aim to assess PackDock’s capability to explore the side-chain conformation space of protein pockets. To reduce the influence of ligand docking algorithms on PackDock, we set ***M*** *and* ***M***^′^ = 1. To balance the docking efficiency and accuracy, we set ***k*** = 1, ***N*** *and* ***N***^′^ = 6, as shown in **Fig. 5(a-d)**.

PackPocket, the core module of PackDock, can generate pocket side-chain conformations for both “free” and ligand “bound” states, as shown in **Fig. 1(b-c)**. Rather than viewing the side-chain prediction task as a regression task, PackPocket defines a diffusion model to treat it as a generative task. This approach provides a better basis for capturing multimodal probability distributions of protein side-chain conformations. The diffusion process uses the side-chain torsion angle score, predicted by a geometric graph neural network at each time step, to denoise a randomly initialized side-chain conformation gradually. The geometric graph neural network consists of three types of nodes: protein residues, protein atoms, and ligand atoms, as shown in **Fig. 1d**. We have defined various types of edges, such as those based on distance, covalent bonds, and relations between protein atoms and protein residues. These edges facilitate information exchange among different nodes, as shown in **Fig. 1e**. The exchanged information is then aggregated to the side-chain atoms, helping to denoise a randomly initialized side-chain conformation, as shown in **Fig. 1f**. Detailed information about our model is provided in **Methods** section.

In summary, PackDock combines conformation selection and induced fit processes by using PackPocket, which allows it to consider different protein-ligand complex binding processes and enhance the efficiency of flexible docking.

### Performance of side-chain prediction in pocket

Protein conformations are not static, especially for the ligand-binding pocket regions, where pocket breathing or allosteric pocket can lead to different side-chain conformations^[14]^. Our model firstly aims to capture the conformation diversity in pocket region. In this section, we mainly discuss the ability of PackPocket to predict accurate pocket conformations while ensuring side-chain diversity. We used two independent test sets: PDBbind time-split test set^[38]^ and the CASP13/14 test set, to evaluate the ability of PackPocket to recover pocket side-chain with or without ligands. Following the training process, we tested side-chain conformations within 5 Å near the ligand for the PDBbind dataset. For the CASP13/14 test set, we extracted the pocket region using the Fpocket^[44]^ pocket-searching tool. We generated the different numbers of pocket conformations by PackPocket, and selected the one closest to the ground truth protein pocket conformation for statistical analysis. For comparison, we also tested several commonly used side-chain packing models/tools, including conventional methods SCWRL^[45]^, FASPR^[46]^ and DL-based methods AttnPacker^[47]^, DiffPack^[48]^. RMSD calculations were performed solely on side-chain atoms from the predefined pocket regions, and the atom accuracy was defined as the fraction of residues having all dihedral angles within 20°of the corresponding native angles.

As shown in **Fig. 2(a-b)**, PackPocket outperformed the current methods on both test sets. PackPocket’s accuracy increased more with higher sampling numbers. At a sampling number of ***N***=40, it achieved 87.8% and 79.1% angle accuracy on the CASP and PDBbind test sets, and atom RMSD was 0.213 Å and 0.336 Å, respectively. We also calculated the atom RMSD and angle accuracy of PackPocket when the sampling number ***N*** was 1, as shown in **Table S3** and **Table S4**. PackPocket still had comparable performance to the current methods. In summary, these experiments confirm PackPocket’s ability to capture accurate pocket conformations while ensuring side-chain diversity and the feasibility of PackPocket to model protein flexibility.

**Fig. 2.**
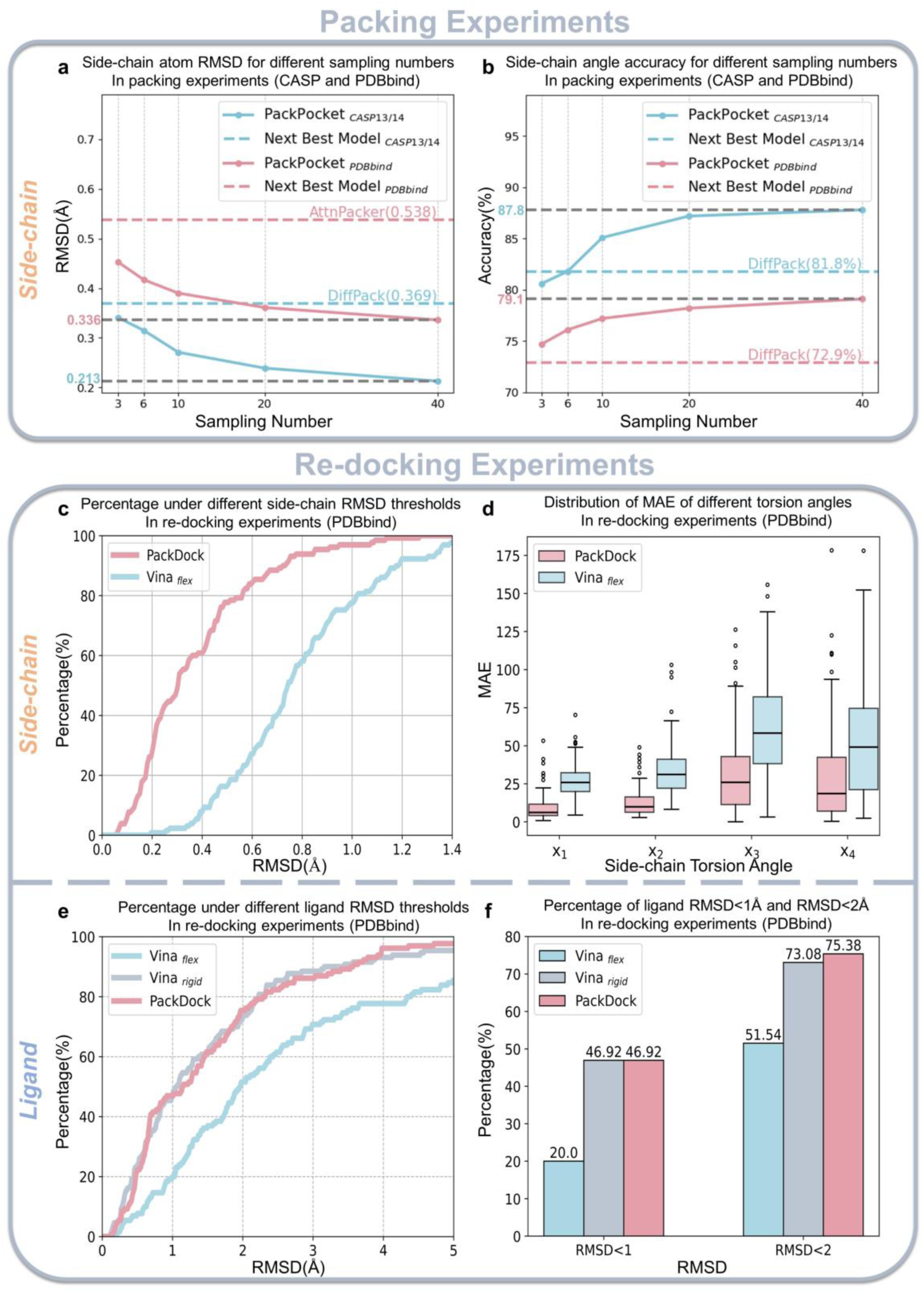
Packing and re-docking experiments across Benchmarks. **(a)** Side-chain atom RMSD for different sampling numbers in packing experiment. Dashed lines indicate the best performance of current methods in packing experiment. **(b)** Side-chain angle accuracy for different sampling numbers in packing experiment. **(c)** Proportion for the flexible docking method under different sidechain RMSD thresholds in re-docking experiment. **(d)** The distribution of MAE for the prediction of side-chain torsion angles by two flexible docking methods in re-docking experiment. **(e)** Proportion for three methods under different ligand RMSD thresholds in re-docking experiment. **(f)** The percentage of ligand RMSD<1 Å and <2 Å for different methods in re-docking experiment.

### Performance of Re-Docking using *holo* structures

To assess the feasibility of PackDock, we mainly discuss the flexible docking performance based on *holo* structures, in which *Vina*_*flex*_ refers to docking with unknown side-chain conformation of *holo*, and *Vina*_*rigid*_ refers to docking with known side-chain conformation of *holo*. It means that rigid docking predicts the ligand binding pose based on the “ground truth” pocket conformation, while flexible docking first predicts the pocket conformation, and then predicts the ligand binding pose based this conformation. Therefore, in this context, the rigid redocking performance potentially represents the upper limit of flexible docking. Moreover, to reduce the influence of the similarity between the proteins in the training set and the test set on the redocking performance, we used the PDBbind time-split test set^[38]^ (no ligand and receptor overlap) for testing.

As shown in **Fig. 2(e-f)**, our method outperformed *Vina*_*flex*_ in redocking, and achieved a success rate similar to *Vina*_*rigid*_ (using *holo* structures). For the percentage of RMSD<1 Å, PackDock’s docking success rate was 47%, which was 26.92% higher than *Vina*_*flex*_’s 20.00% success rate. For the percentage of RMSD<2 Å, PackDock’s success rate reached 75.38%, which was 23.84% higher than *Vina*_*flex*_. We further analyzed the side-chain conformation generated by the two flexible docking methods. As shown in **Fig. 2c**, the RMSD of the side-chain conformation generated by PackDock was significantly outperformed than that of *Vina*_*flex*_. We also performed a statistical analysis of the mean absolute error (MAE) of different torsion angles of the side chains, and our method achieved a significant predict accuracy improvement for each torsion angle compared to *Vina*_*flex*_, as shown in **Fig. 2d**. Overall, due to the high accuracy of PackPocket’s side-chain prediction, PackDock’s flexible docking performance was similar with the rigid docking using the *holo* structures, and significantly surpassed *Vina*_*flex*_. In summary, these redocking experiments demonstrated the ability of PackPocket to accurately predict side-chain conformations and the feasibility of PackDock’s docking process.

### Performance of Cross-Docking using *apo* structures

In most real-world application situations, it is challenging to obtain the *holo* structure that binds with the ligand of interest. Often, we can only use *apo* or non-homologous ligand-induced *holo* structures for docking simulation. In these cases, treating the protein as rigid and only considering the flexibility of the ligand may have significant limitations. This section aligns with these real-world scenarios by using *apo* structures. Previously, Zhang et al.^[49]^ collected 32 targets‘ *apo* from the DUD-E database^[50]^, each with multiple *holo* structures bound with different non-homologous ligands, and categorized into three groups based on the α-C RMSD at the ligand binding site: group 1 with α-C RMSD <0.5 Å, group 2 with 0.5 Å < α-C RMSD < 1.5 Å, and group 3 with α-C RMSD > 1.5 Å. We used this dataset to test the cross-docking performance of *apo* structures. For each method, we generated a total of 36 complex conformations and selected the one closest to the ground truth ligand binding pose for subsequent statistical analysis. Based on the previous work^[51]^, we define the flexible region as side chains that are within 3.5 Å from the ligand for all the conventional flexible docking algorithms, including: *Autodock*_*flex*_ ^[52]^, *Vina*_*flex*_^[34]^, *Smina*_*flex*_^[53]^, and *Gnina*_*flex*_^[54]^.

**Fig. 3(a-b)** shows that PackDock significantly outperforms existing docking methods across all targets. **Table.1** further illustrates the average RMSD of ligands in different groups and the percentage of ligand RMSD < 1 Å or < 2 Å for various methods: in group 1, PackDock’s average ligand RMSD of 1.73 Å, with 70.83% of RMSD being less than 2 Å and 31.25% of RMSD being less than 1 Å. The best among other methods is Diffdock, with an average ligand RMSD of 2.14 Å, 68.75% of RMSD below 2 Å, but only 10.41% of RMSD below 1 Å. In group 2, PackDock’s average ligand RMSD of 2.52 Å, while the next best method, *Autodock*_*flex*_, has only 3.31 Å. PackDock also has 47.50% of RMSD being less than 2 Å and 16.25% of RMSD being less than 1 Å, surpassing the current methods by 18.75% and 11.25%, respectively. For group 3, which has a large α-CRMSD, most methods did not achieve satisfactory docking accuracy, but PackDock still had an average ligand RMSD of 2.86 Å and 30.92% of ligand RMSD being less than 2 Å. Overall, in group 2 and 3, as the backbone RMSD increases, all methods’ performance slightly decreases. However, our method still maintains a high performance. In all groups, PackDock increased the docking success rate by 12.50% compared to all other methods. We also compared the re-docking experiments using *holo* structures and the cross-docking experiments using *apo* structures. Although PackDock and *Vina*_*rigid*_ have similar performance in re-docking when using *holo* structures, PackDock shows a 21.43% improvement over *Vina*_*rigid*_ in docking performance when using *apo* structures. This underscores the limitations of rigid docking and the importance of considering flexibility of side chains. In conclusion, PackDock can effectively consider ligand-induced protein conformational changes, and accurately predict the binding pose of ligands based on *apo* structures.

**Fig. 3.**
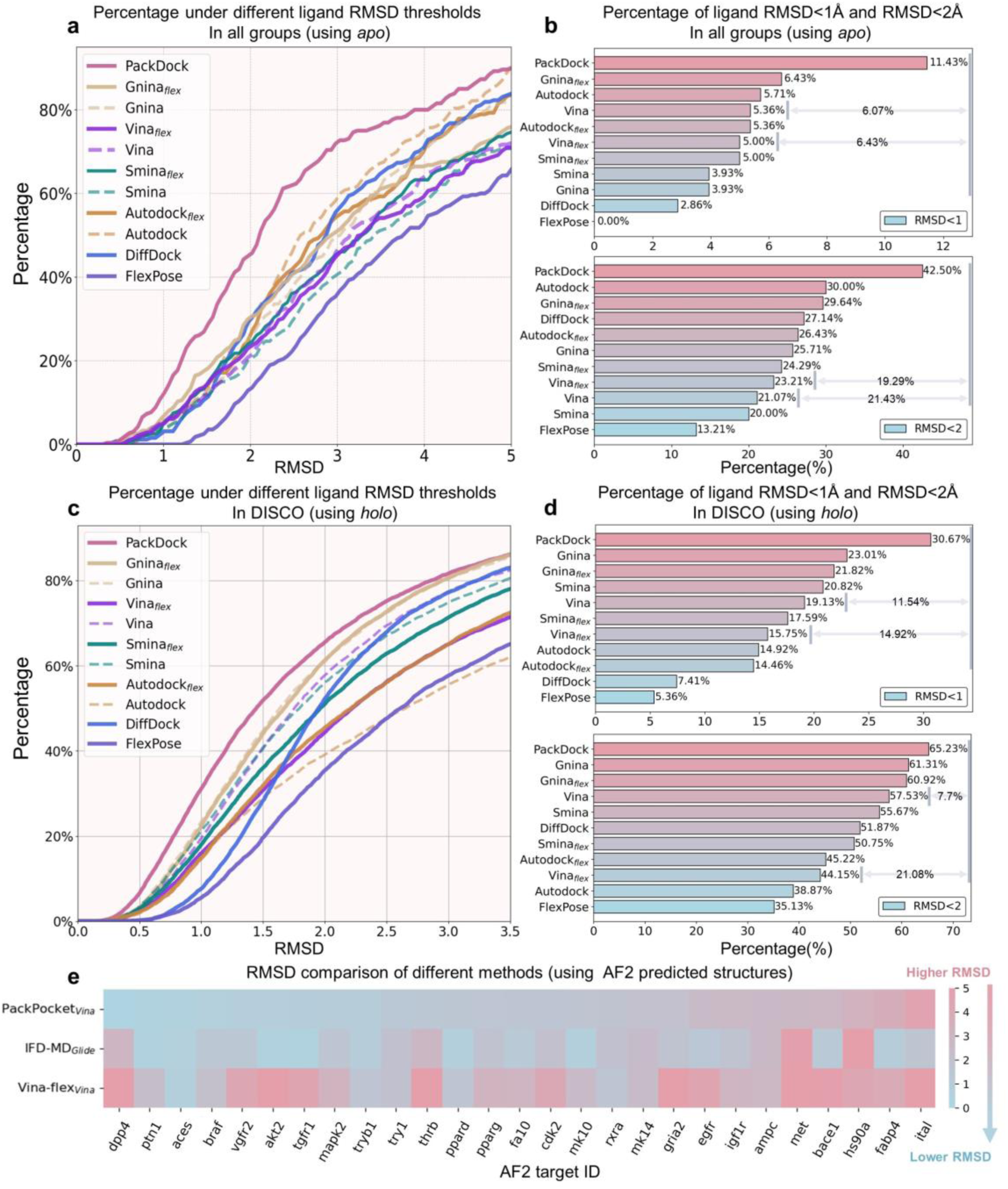
Cross-Docking experiments across Benchmarks. | **(a)**The percentage of ligand RMSD under different thresholds in all groups, when using *apo*. **(b)** The percentage of ligand RMSD<1 Å and <2 Å for different methods in all groups, when using *apo*. **(c)**The percentage of ligand RMSD under different thresholds in DISCO, when using *holo*. **(d)** The percentage of ligand RMSD<1 Å and <2 Å for different methods in DISCO, when using *holo*. **(e)** The ligand RMSD of docking with the predicted structure by AF2.

**Table.1.** Cross-Docking experiments with *apo* structure. Shown is the percentage of predictions with RMSD < 2Å and RMSD < 1 Å, and the mean RMSD for all groups and the three groups separately. The last line shows our method’s performance. Best results are in bold and second best are underlined in two categories.

| Method | Group1 |  |  | Group2 |  |  | Group3 |  |  | All Groups |  |  |
| --- | --- | --- | --- | --- | --- | --- | --- | --- | --- | --- | --- | --- |
|  | %<1 Å | %<2 Å | Mean | %<1 Å | %<2 Å | Mean | %<1 Å | %<2 Å | Mean | %<1 Å | %<2 Å | Mean |
| *Autodock* | <u>18.75</u> | 43.75 | 2.47 | 3.75 | 20.00 | 3.33 | 2.63 | **30.92** | <u>2.96</u> | 5.71 | <u>30.00</u> | <u>2.98</u> |
| *Smina* | 2.08 | 29.16 | 2.91 | 1.25 | 16.25 | 5.21 | **5.92** | 19.07 | 3.62 | 3.93 | 20.00 | 3.95 |
| *Gnina* | 6.25 | 39.58 | 2.75 | 1.25 | 20.00 | 3.88 | <u>4.61</u> | 24.34 | 3.19 | 3.93 | 25.71 | 3.31 |
| *Vina* | 8.33 | 29.16 | 2.83 | 1.25 | 15.00 | 4.38 | **5.92** | 21.71 | 3.61 | 5.36 | 21.07 | 3.70 |
| *Diffdock* | 10.41 | <u>68.75</u> | <u>2.14</u> | 1.25 | <u>28.75</u> | 3.35 | 1.31 | 13.15 | 3.67 | 2.86 | 27.14 | 3.28 |
| *FlexPose* | 0.00 | 12.50 | 4.06 | 0.00 | 18.75 | 3.96 | 0.00 | 10.52 | 4.39 | 0.00 | 13.21 | 4.21 |
| *Autodock<sub>flex</sub>* | 16.66 | 35.41 | 2.71 | <u>5.00</u> | 27.50 | <u>3.31</u> | 1.97 | 23.02 | 3.33 | 5.36 | 26.43 | 3.22 |
| *Smina<sub>flex</sub>* | 4.16 | 43.75 | 2.66 | 3.75 | 16.25 | 4.13 | **5.92** | 22.36 | 3.59 | 5.00 | 24.29 | 3.58 |
| *Gnina<sub>flex</sub>* | 14.58 | 47.91 | 2.43 | <u>5.00</u> | 26.25 | 3.65 | <u>4.61</u> | 25.65 | 3.60 | <u>6.43</u> | 29.64 | 3.41 |
| *Vina<sub>flex</sub>* | 10.41 | 37.50 | 2.78 | 3.75 | 16.25 | 4.46 | <u>4.61</u> | 22.36 | 3.73 | 5.00 | 23.21 | 3.78 |
| PackDock | **31.25** | **70.83** | **1.73** | **16.25** | **47.50** | **2.52** | 2.63 | **30.92** | **2.86** | **11.43** | **42.50** | **2.55** |

### Performance of Cross-Docking using *holo* structures

Treating protein structures induced by non-homologous ligands as rigid could result in inaccurate ligand binding poses. This section primarily discusses flexible docking tests when only non-homologous ligands’ protein structures are available. DISCO ^[55]^ is a large-scale Cross-Docking test set. Given the complexity of cross-docking, previous work down sampled DISCO to obtain 7970 protein-ligand structure pairs^[54]^, where no receptor was paired with its homologous ligand. These 7970 protein-ligand structure pairs were used as the Cross-Docking test set in this study. We generated a total of 36 complex conformations for each method and selected the one closest to the ground truth ligand binding pose for statistical analysis and define the flexible region as side chains that are within 3.5 Å from the ligand for all the conventional flexible docking algorithms.

In **Fig. 3c**, the proportion of ligand poses falling below different RMSD thresholds were visualized, and PackDock’s curve is higher than the rest of the methods. Then, we separately counted the docking success rates under different RMSD threshold values, as shown in **Fig. 3d**. PackDock achieved 30.67% and 65.23% success rates for RMSD<1Å and RMSD < 2Å, respectively. The second-best method, Gnina, had 23.01% and 61.31% docking success rates, respectively. For the more stringent criterion of RMSD < 1Å, PackDock significantly improved the docking success rate, while DL-based methods, Diffdock and FlexPose had much lower success rates. This result is consistent with the observation of Cross-Docking using *apo* structures in **Fig. 3b**. It should be noted that the conventional flexible docking algorithms did not show more obvious advantages than the rigid docking methods in most cases. This could be because the same protein typically does not alter much in the binding pocket when interacting with different ligands, As shown in Supplementary **Fig. S1** and **Fig. S2**, most *holo* structures of the same protein have RMSD values less than 2 Å. Similar observations were also made in previous studies ^[54]^, where they found that flexible docking might increase the ligand RMSD compared to rigid docking when the binding pocket similarity was high. This is consistent with the conclusions from the redocking tests: while conventional algorithms may increase the diversity of the side chain conformation, they typically struggle to accurately identify those near the native conformations. In contrast, PackPocket has a significant advantage in the accuracy of side-chain prediction, and PackDock can more comprehensively consider the side-chain conformation space by combining both conformation selection and induced fit processes. This enables PackDock to have a high ligand prediction accuracy when the pocket similarity is high or low. In summary, through testing on the Cross-Docking dataset, PackDock has demonstrated its significant advantages over conventional flexible docking methods and DL-based docking methods. It has also proven the feasibility of flexible docking when only non-homologous ligands’ protein structures are available.

### Performance of Cross-Docking using AF2 predicted structures

Experimental methods have determined about 100,000 structures over the years^[56]^, which only cover a small part of the known protein sequence space^[57]^. Homology modeling or AI-based protein folding algorithms can be used as alternatives to obtain protein structures in an efficiency manner. However, the docking accuracy based on the protein structures obtained by computational methods is poor due to the insufficient consideration of side chain packing^[26]^. Previous studies have attempted to mitigate this issue using physics-based methods, such as the IFD-MD^[58]^. In this section, we use the AF2 predicted structure as an example to evaluate whether PackPocket can enhance the docking performance of the computationally predicted structure.

In this docking experiment, we used the dataset of Zhang et al.^[58]^ as the test set, which includes the *holo* and AF2 predicted structures of 27 targets. This dataset also contains docking result based on the IFD-MD refined AF2 predicted structures^[58]^, which were obtained by using the aligned crystal ligand pose as the template to induce the protein conformation. For comparison, we followed the same procedure to use PackDock to generate pocket conformation. For PackPocket and *Vina*_*flex*_, a total of 36 complex conformations for each method were generated and the one closest to the ground truth ligand binding pose was selected for statistical analysis. As results shown in **Fig. 3e**, PackPocket + *Vina* can correctly identify the binding mode for most of the targets. Additionally, the average RMSD (1.77Å) of PackPocket significantly outperformed *Vina*_*flex*_ (3.68Å) and showed comparable performance to IFD-MD + *Glide* (1.81Å). In summary, this section primarily discusses the performance of using AF2 predicted structures in docking experiments. It demonstrates that PackPocket can enhance the performance of AF2 predicted structures in related tasks. This is of great significance for the SBDD community, as it provides a feasible approach for SBDD starting from sequences.

### Identification of Key Amino Acid Conformational Distribution

To delve deeply into the performance of PackDock in predicting side-chain conformations, we present a series case studies. In these cases, we start with *apo*, aiming to predict *holo* structure after ligand binding. **Fig. 4** displays the multiple side-chain conformations predicted by PackDock, and provides the corresponding *apo*, *holo*, and ground-truth ligand pose as references for comparison. The docking pipeline accurately identified the amino acid residues that differed substantially between *apo* and *holo*, and also revealed the possible conformational distribution preferences. As shown in **Fig. 4a**, in the *apo* (5e8s, green)-*holo* (3hmm, cyan) structure, only D351 has a significant side-chain conformation difference. PackDock successfully sampled the side-chain conformation distribution of this amino acid, and identified the conformation that matches the *holo* structure. It also captured the bimodal conformation distribution of this amino acid at this site (consistent with the *apo* and *holo*). For the rest of the amino acids, there are only slight conformational differences, and PackDock also generated compact unimodal conformation distribution for them. Similarly, as shown in **Fig. 4b**, F31 in *apo* (1pdb, green) and *holo* (1boz, cyan) shows significant differences in the key amino acid conformations, and PackDock accurately captured these conformation shift. The same phenomenon was also observed in other systems (**Fig. S3**). We further analyzed the torsion angle distribution generated by PackDock, as shown in **Fig. 4(c-d)**. For amino acids with substantial variations in side-chain torsional angles between *apo* and *holo* states, PackDock can sample a broad range of torsional angle distributions, as observed in D351 in *apo*(5e8s)-*holo*(3hmm) and F31 in *apo*(1pdb)-*holo*(1boz). In contrast, for amino acids with minimal changes in side-chain torsional angles between *apo* and *holo* structures, PackDock samples more concentrated torsional angle distributions, exemplified by Y282 and H283 in *apo*(5e8s)-*holo*(3hmm) and F34 and Y121 in *apo*(1pdb)-*holo*(1boz). This demonstrates that the side-chain conformations generated by PackDock are accurate and representative. This advantage can reduce the complexity of the flexible docking process, thereby increasing the success rate of docking, which is consistent with the results of redocking. Furthermore, it can accurately identify the key amino acid sites in *apo* related to the ligand. This can provide a feasible direction for the optimization of the lead compound stage, which is of great significance for achieving more precise SBDD.

**Fig. 4.**
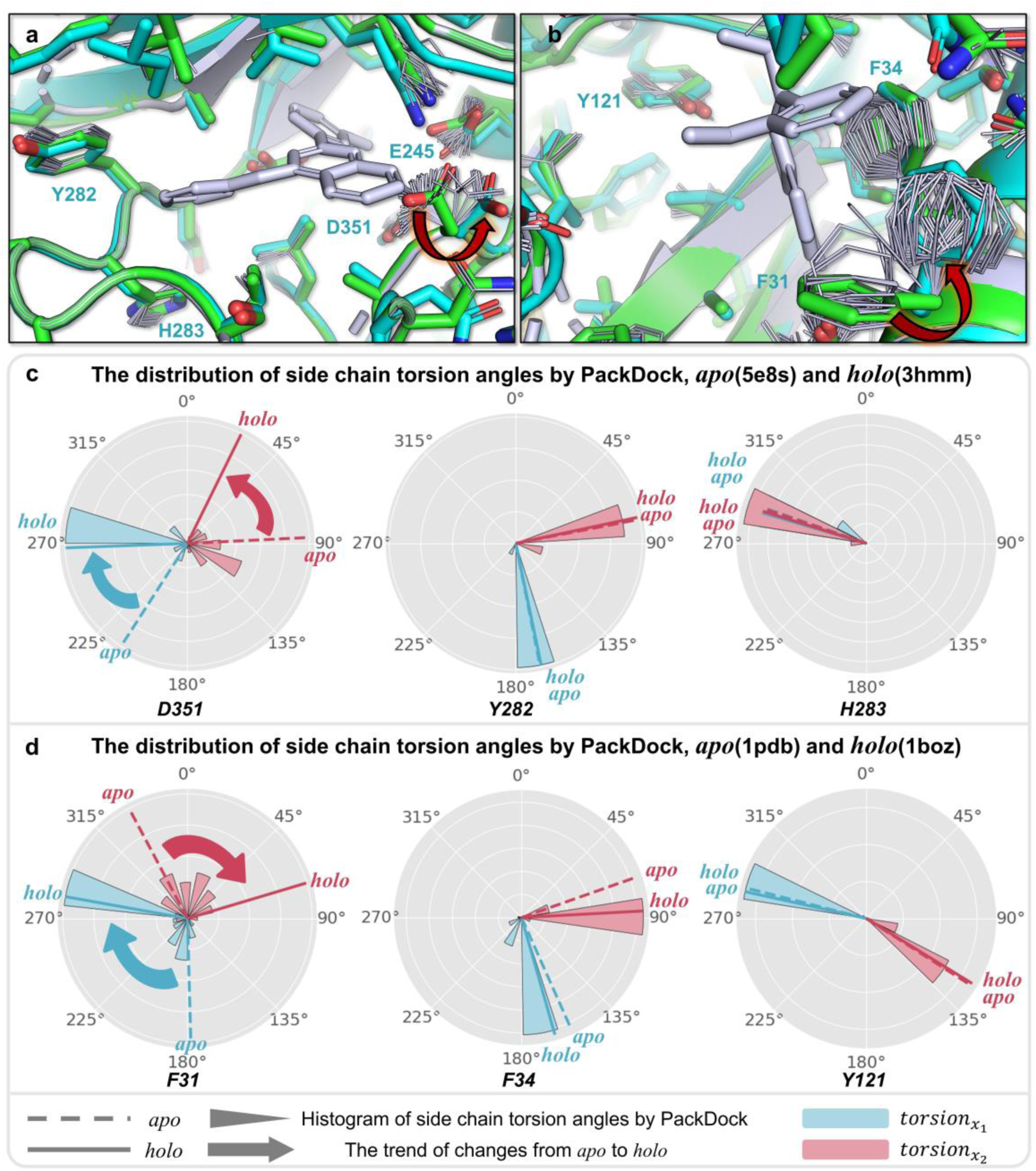
PackDock captured conformational trends. **(a)** In the *apo* (5e8s, green) – *holo* (3hmm, cyan) structure, D351 exhibits a significant conformational change. **(b)** In the *apo* (1pdb, green) - *holo* (1boz, cyan) structure, F31 exhibits a significant conformational change. Other amino acids with smaller conformational differences show a compact unimodal distribution. **(c)** The statistics of the side-chain torsion angles of D351, Y282, and H283 generated by PackDock for *apo*(5e8s)-*holo*(3hmm) **(d)** The statistics of the side-chain torsion angles of F31, F34, and Y121 generated by PackDock for *apo*(1pdb)-*holo*(1boz)

### Docking runtime evaluation

One of the challenges of flexible docking methods is the increased computational complexity. In PackDock, as shown in **Fig. 1a**, there are several hyperparameters that affect the running time and docking success rate, such as the number of docking ligand conformations ***M*** and ***M′***, the number of side-chain sampling ***N*** and ***N′***, and the number of iterations for the induced-fit stage ***k***. This section analyzes how the number of complex samples and the iterations of the induced-fit stage affect the running time and docking success rate of PackDock, and to reduce the impact of different docking algorithms on PackDock, we fix ***M*** and ***M′*** to 1. We first experimented with the sampling number of PackDock, and as shown in **Fig. 5(a-b)**, we found a linear relationship between the sampling number and the running time, and the optimal docking performance was achieved with 36 samples (***N*** *and* ***N***^′^ = 6, ***k*** = 1). Then we analyzed the iteration number ***k*** of the induced-fit stage, and as shown in **Fig. 5(c-d)**, the docking success rate depends largely on whether the induced-fit stage is performed or not. However, further increasing the number of the induced-fit stage did not improve the docking performance significantly. Overall, by combining the conformation selection stage and the one-step induced-fit stage, PackDock achieved a balance between the docking performance and the computational time. When keeping a similar search space size (setting the side chain flexibility within 5 Å of the ligand), our method was nearly 10 times and 25 times faster than SMINA and IFD, respectively, as shown in **Fig. 5(e-f)**. In addition, by analyzing the distribution of the docking time, we found that most of the time was allocated to the serial docking. This implied the prospect of improving the work efficiency by using parallel computing in the docking stages.

**Fig. 5.**
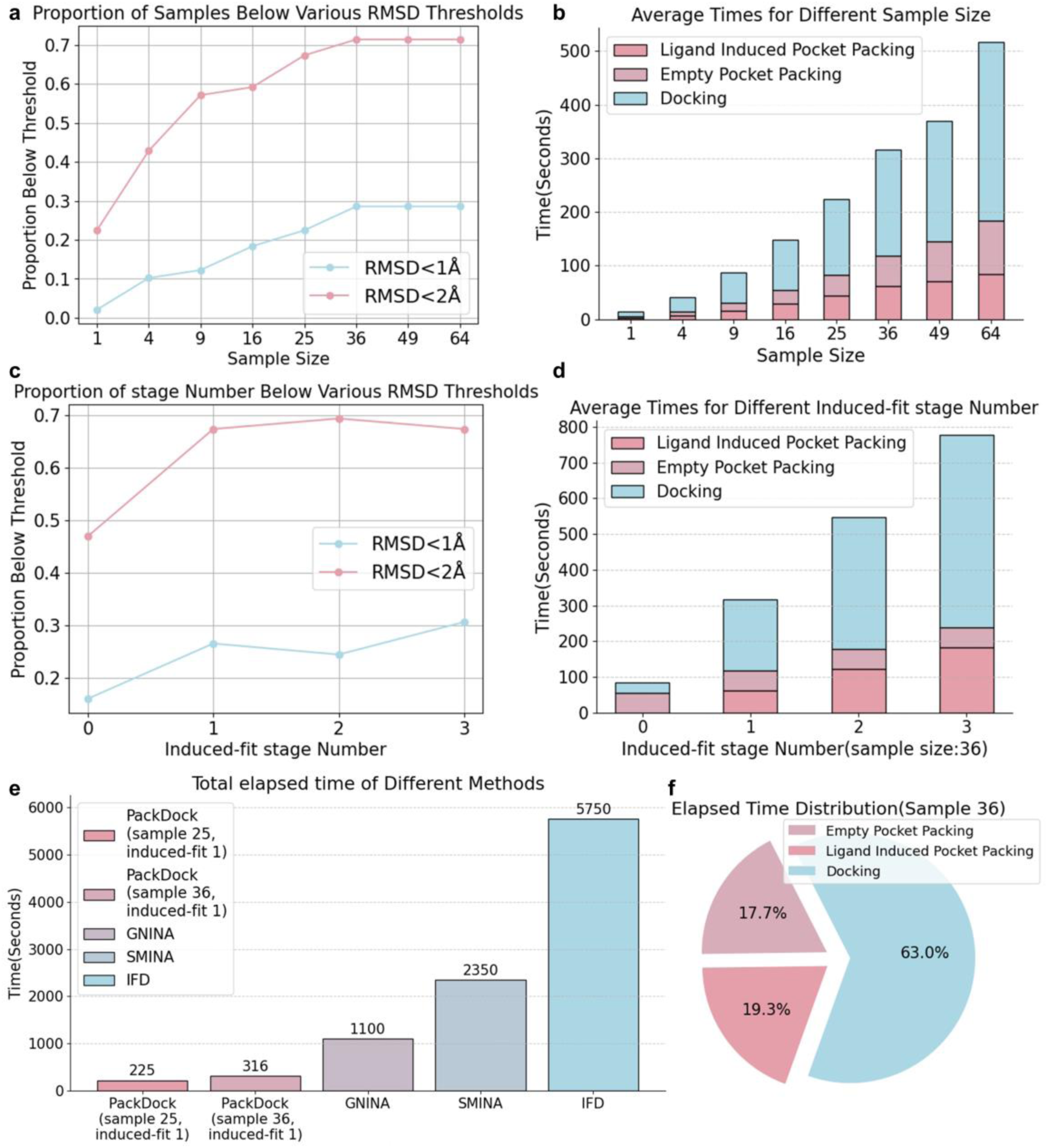
Running time analysis. **(a)** Relationship between different sampling number and docking accuracy. **(b)** Relationship between different sampling number and docking time. **(c)** Relationship between different induced fit cycle numbers and docking accuracy. **(d)** Relationship between different induced fit cycle numbers and docking time. **(e)** Comparison of docking time with traditional flexible methods. **(f)** Distribution of docking time with default PackDock settings.

## Discussion

The rapid advancement of structural biology and AI-based protein folding algorithms has enabled access to the protein structures of more novel drug targets. However, these protein structures often don’t meet the needs of drug design due to the lack consideration of flexibility upon ligand binding. This study proposes a new flexible docking solution, PackDock, to addresses the limitations of existing static structures for drug design. Specifically, we leverage the diffusion model’s generative diversity and conditional generation capabilities to simulate protein pocket conformations in either the ligand “free” state or the ligand “bound” state. Based on the diffusion model, we established a flexible docking pipeline, which combines conformational selection theory and induced-fit theory to account for different binding processes of protein-ligand complex. We’ve designed various tests to assess PackDock’s generalization capability to different initial static protein or protein-ligand complex structures: (1) The side-chain recovery tests suggested that it could accurately predict the side-chain conformations of the ligand binding regions. The re-docking tests verified the feasibility of PackDock from both the side-chain and the ligand perspectives. (2) The cross-docking tests with apo and non-homologous ligand-induced holo demonstrated PackDock’s ability to accurately predict the protein-ligand complex structure in the most common SBDD scenarios, proving its high practical value. (3) The AF2 test verified that PackPocket could enhance the docking performance based on hypothetical structural model. Overall, PackDock is capable of processing various protein structures and predicting the structures of protein-ligand complexes with high accuracy, expanding the applicability domain of SBDD.

The mechanisms of biomolecular recognition are closely linked to proteins’ intrinsic dynamics. It remains a challenge to effectively account for protein flexibility upon ligand binding. PackDock employs a data-driven method to address this longstanding problem, using deep generative learning from existing static structure data to gain insights into the packing preferences of amino acid side chains. This knowledge is crucial for understanding molecular interactions and the dynamic behavior of proteins. Combining the recent advances in cryo-electron microscopy and protein structure prediction, we envisage that PackDock will become an essential tool of the community, and provide a more realistic perspective for understanding ligand-protein binding process.

## METHODS

### Model Overview

PackPocket focuses on modeling the side chains of ligand binding sites. A protein side-chain conformation represents a specific arrangement of side-chain atoms in three-dimensional space, we can regard the protein side-chain conformation *x* as an element in ℝ^3×*n*^, where *n* is the number of atoms. Due to the predominantly rigid nature of bond lengths, angles, and small rings within side-chain conformations, the flexibility of the side chain primarily resides in the torsional angles of rotatable bonds. Hence, the conformational space consistent with the initial protein side-chain conformation *x* forms an *m*-dimensional submanifold *Mx* ∈ ℝ^*m*^, where *m* represents the number of rotatable bonds within the side-chain. Side-chain packing is defined as learning the probability distribution *p*(*x*) concerning the manifold *Mx*. Simultaneously, PackPocket defines a unique way (Supplementary **Table S1**) to determine the side-chain torsion angles of proteins, ensuring a one-to-one mapping between the torsion space and the three-dimensional conformation space. To facilitate the model learning the torsion angle distribution, we defined the rotation direction of atoms at both ends of a torsional angle by rotating only the section near the terminal end of the side chain, which also kept the coordinates of the main chain atoms unchanged during the sampling process. During the sampling process, PackPocket excludes Proline residues from sampling and prevents the breakage of disulfide bonds between Cysteine residues. Additionally, following pervious works ^[41, 59, 60]^, we define the score matching objective function in torsional space but handle the conformations directly in three-dimensional coordinates.

### Score Model

To train a score-based diffusion model on protein side-chain space, we defined a bijective side-chain torsion angle definition (Supplementary **Table S1**) and constructed a scoring model, *s*(*x*, *l*; *t*), that takes as input the three-dimensional structure of the protein conformation *x*, the pose of the ligand pose *l* (if exist), and the diffusion time coding *t*. The network outputs torsion angle scores invariant under SE(3) transformations for rotatable side-chain bonds near the protein pocket. Score model is constructed using the e3nn library^[61]^.

Model details are depicted in **Fig. 1**, where the scoring network is structured in the form of a geometric heterogeneous graph. The nodes include the protein pocket heavy atoms and ligand heavy atoms (if exist). When constructing edges, in addition to considering distance edges and covalent edges between nodes, connections are established solely between the protein heavy atom nodes and their corresponding residue nodes. To streamline the complexity of distance edges and enhance efficiency, we applied distinct cutoff values to different node types. For example, we applied a cutoff of 5 Å between atomic nodes, 30 Å between residue nodes, and a maximum of 24 neighbors per node.

The residue nodes utilize the amino acid type as initial features, while receptor heavy atom nodes are assigned predefined initial features, and ligand atoms’ features are computed using RDKit and encompass: atomic number; stereochemistry; degree; formal charge; implicit valence; number of attached hydrogen atoms; number of free valence electrons; hybridization type; and whether the atom is in an aromatic ring, etc. These features are concatenated with the sine positional encoding for diffusion time ^[62]^ and edge features used radial basis function encoding for edge lengths *r*_*ab*_^[63]^. These scalar features of each node and edge are transformed into a set of scalar features through a learnable MLP for initial representations as interaction layers (Each node and edge type uses a parameter-independent MLP).

In the Embedding update layer, we utilized initial representations of different nodes and constructed information exchange between nodes using spherical harmonics of edge vectors through tensor products. Subsequently, each node aggregated information from connected nodes via various edges. We updated node features through residual connections, as depicted in equation (1), achieving the objective of feature aggregate:

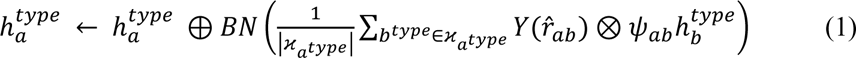

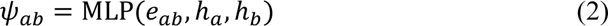

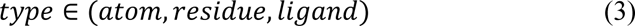

where ℎ_*a*_ and *e*_*ab*_ represent the representation of node *a* and edge *ab*, respectively. ⊕ denotes residual connections, *BN* signifies equivariant batch normalization, *ϰ*_*a*_ refers to neighboring nodes of *a*, *Y* represents spherical harmonics functions with ℓ maximum of 2, ⊗ denotes the tensor product of irreps representations of spherical harmonics^[61]^. Equation (3) shows the different types of nodes. Distinct convolutional layers were employed for nodes of different types, as illustrated in **Fig. 1**. After embedding all types of atoms, we identified the rotatable side chains and embedded their terminal and nearby atoms.

To predict the m SE(3)-invariant torsion scores, we built an additional convolutional layer, inspired by previous work [41,61], that focused on the “rotatable bonds”. Specifically, for each “rotatable bond” *g* = (*g*_0_, *g*_1_) and *b* ∈ V, we obtain the node embedding ℎ and the radial basis embedding *μ*(*r*) of the edge length, as shown in equations (4), (5), and (6). Then, we constructed a convolutional filter *T*_*g*_ for each bond *g* from the tensor product of the spherical harmonics with ℓ = 2 representation of the bond axis 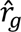, as depicted in equation (7). Next, we used the filter *T*_*g*_ to perform convolution on the neighboring nodes of the bond, obtaining representations for the “rotatable bond”, as in equation (8).

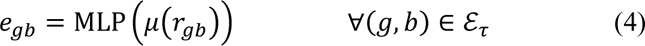

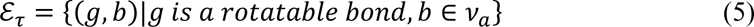

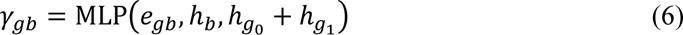

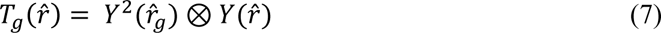

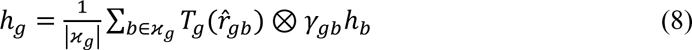

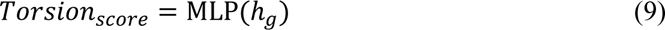

Finally, a MLP is employed to predict the final torsion angle scores *Torsion*_*score*_, as equation (9). The parameters of each MLP are not shared.

### Dataset construction

To evaluate the accuracy of PackPocket in predicting the side chains of pocket regions under different scenarios, we used the protein dataset BC40^[64]^ and the protein-ligand complex dataset PDBbind ^[65]^.

BC40 represents a subset derived from the PDB database, curated by filtering sequences with a 40% similarity threshold, resulting in 36,000 proteins. To focus more on modeling side-chain conformations near the binding pockets, we employed the Fpocket tool^[44]^ to identify potential pockets within these proteins, discovering approximately 1.5M pocket regions. Next, we obtained about 38,000 pocket regions by ranking and scoring with Fpocket. Finally, we defined the pocket graph as the amino acids within 12 Å radius from the pocket region center, and then trained the model on these graphs to predict the side chain conformation in the empty pocket. The dataset was randomly split and tested using external test sets CASP 13 and 14. The PDBbind v2020 dataset contains 19,443 protein-ligand complexes, covering 3,890 unique receptors and 15,193 unique ligands. We excluded all complexes that the RDKit couldn’t handle, ultimately selecting 19,119 complexes for training a side-chain conformation prediction model based on ligand conformations. Additionally, following previous work^[38]^, we split the dataset into training, validation, and test sets according to the release time of the complex structures. For the PDBind dataset, we defined the pocket graph as the amino acids that have at least one atom within 8 Å of the ligand position, and then trained the ligand induced pocket packing model on these graphs. Due to the abundance of hydrogen atoms in proteins and ligands, we used Rdkit’s implicit hydrogen representation to construct the pocket graph. All training data underwent processing through OpenBabel.

### Training details

We trained the empty pocket packing model and the ligand induced pocket packing model using the Variance-Exploding SDE (VE-SDE) framework for training. The function *σ*(*t*) is chosen to follow the exponential decay defined in previous research^[66]^, as *σ*(*t*) = (0.01*π*)^1−*t*^(*π*)^*t*^, *t* ∈ (0, 1). During training, the protein side-chain conformation is randomly perturbed by diffusion kernels at each time step *t*, and then the score model takes the noisy conformation as input to estimate the noise. We used Adam ^[44]^ as the optimizer for the score model. We updated the moving average with a decay factor of 0.999, learning rate of 1e-3, and batch size of 8. Every 5 epochs, we ran the inference step to infer on the validation set, and used the epoch with the smallest average side chain RMSD as the final diffusion model. Two models were trained on two 80GB A100 GPUs for 200 epochs, taking a total of about 20 days.

### Docking Pipeline

For stage 1, we first generate 36 pocket conformations using the empty pocket packing model given the pocket backbone conformation, then calculate the RMSD matrix and cluster them into 6 groups using the K-Medoids algorithm, and then use the medoids of each cluster as the representative conformation, thus simulating the conformation selection theory. After obtaining 6 pocket conformations similar to the “free” state, we dock one ligand conformation for each of them using *Vina*-GPU^[67]^ (for work efficiency, the docking process uniformly uses GPU-accelerated *Vina*). For stage 2, we use the ligand induced pocket packing model with the 6 ligand conformations obtained in the stage 1 as the initial condition. Then, for each docked ligand pose, we generate 6 corresponding protein pocket conformations by conditional generation. This produces 36 induced pocket conformations. Finally, we obtain 36 complete protein-ligand complex conformations as output by docking. To consider the environmental information around the pocket and the ligand-induced effect, we select the flexible amino acids for packing in the empty pocket packing model and the ligand induced pocket packing model, respectively. These amino acids are within 0.8 * volume of the pocket region and 5 Å around the ligand.

### Evaluation Metrics

#### Angle MAE, Angle Accuracy, Side-chain RMSD

We assess the accuracy of generated side-chain conformations using three metrics: (1) Angle MAE, which quantifies the mean absolute error in predicted torsional angles. (2) Angle Accuracy, indicating the percentage of correct predictions, with a torsional angle considered accurate if the deviation is within 20 degrees. (3) Side-chain RMSD, measuring the average root mean square deviation of side-chain atoms for each residue within 5 Å of the ligand and near the pocket center within 8 Å.

#### Ligand RMSD, Docking success rate

The ligand RMSD values between generated ligand binding poses and crystallized ligand binding poses were calculated by the obrms module of OpenBabel. The docking accuracy was measured by the success rate of generating binding conformations with the RMSD of less than 2 Å from the ground-truth conformations.

## Supporting information

PackDock-Supplementary-Information

## Data availability

The data included in our paper are all from public data sets.

## Code availability

The code used to generate the results shown in this study will be available under an MIT License in the repository https://github.com/Zhang-Runze/PackDock upon publication.

## ACKNOWLEDGMENTS

We gratefully acknowledge financial support from National Key Research and Development Program of China (2022YFC3400504 and 2023YFC2305904), National Natural Science Foundation of China (T2225002 and 82273855), SIMM-SHUTCM Traditional Chinese Medicine Innovation Joint Research Program (E2G805H), the open fund of state key laboratory of Pharmaceutical Biotechnology, Nanjing University, China (KF-202301).

## Notes

### Competing Interest Statement

The authors have declared no competing interest.

