## Supplementary material for "PackDock: a Diffusion Based Side Chain Packing Model for Flexible Protein-Ligand Docking": PackDock-Supplementary-Information

Runze Zhang[▽,1,2], Xinyu Jiang[▽,1,2], Duanhua Cao[▽,1,3], Jie Yu[1,4,5], Mingan Chen[1,4,6], Zhehuan Fan[1,2], Xiangtai Kong[1,2], Jiacheng Xiong[1,2], Zimei Zhang[1,7], Wei Zhang[1,2], Shengkun Ni[1,2], Yitian Wang[1,2], Shenghua Gao[5], Mingyue Zheng[*,1,2]

[1] Drug Discovery and Design Center, State Key Laboratory of Drug Research, Shanghai Institute of Materia Medica, Chinese Academy of Sciences, 555 Zuchongzhi Road, Shanghai 201203, China

[2] University of Chinese Academy of Sciences, No. 19A Yuquan Road, Beijing 100049, China

[3] Innovation Institute for Artificial Intelligence in Medicine of Zhejiang University, College of Pharmaceutical Sciences, Zhejiang University, Hangzhou, Zhejiang 310058, China

[4] Lingang Laboratory, Shanghai, 200031, China

[5] School of Information Science and Technology, ShanghaiTech University, 393 Middle Huaxia Road, Shanghai 201210, China

[6] School of Physical Science and Technology, Shanghai Tech University, 393 Middle Huaxia Road, Shanghai, 201210, China

[7] Division of Life Science and Medicine, University of Science and Technology of China, Hefei, Anhui, 230026, China

**Table of Contents**

S1 Baseline Method

S2 The Definition of Side-chain Torsion Angles

S3 Packing Dataset

S4 Performance of Packing Experiments

S5 Pocket Change Statistical and Experiment Number Statistical in Cross-Docking

S6 Visualization of Side-chains in PackDock

### S1. Baseline Method

#### S1.1 Packing Method

SCWRL4[45]. SCWRL4 is a widely-used method for predicting protein side-chain conformations. It employs a graph-based approach, treating side chains as nodes connected by edges representing potential steric clashes. The algorithm searches for the most probable side-chain conformation, minimizing clashes and optimizing energy using a backbone-dependent rotamer library and a statistical potential energy function based on known protein structures.

FASPR[46]. FASPR (Fast and Accurate Side-chain Prediction using Rotamer libraries) efficiently predicts side-chain conformations by utilizing backbone-dependent rotamer libraries and a custom energy function. The method employs Dead-End Elimination (DEE) and tree decomposition to identify the rotamers leading to the Global Minimum Energy Conformation (GMEC).

AttnPacker[47]. AttnPacker is a DL-based method for side-chain conformation prediction that utilizes the attention mechanism, a powerful technique commonly employed in deep learning architectures for tasks involving sequence data. The protocol and settings in notebook *protein_learning/examples/inference.ipynb* in the AttnPacker repository ( https://github.com/MattMcPartlon/AttnPacker ) were used for packing.

DiffPack[48]. DiffPack is a torsional diffusion model designed for predicting the conformation of protein side-chains based on their backbones. The protocol and settings in *script/inference.py* in the DiffPack repository ( https://github.com/DeepGraphLearning/DiffPack ) were used for packing.

#### S1.2 Docking Method

*Autodock*[52] The software versions employed include *Autodock*-GPU, Reduce 3.23.130521, ADFRsuite 1.0, and RDKit 2022.09.1. Ligands are processed into PDBQT files using ADFR *prepare_ligand* scripts, and RDKit calculates their centroid coordinates and coordinate ranges. Hydrogen atoms are added with Reduce, and

PDBQT files are generated using the ADFR *prepare_receptor script*. The docking box size is set using the co-crystallized ligand's center and its maximum diameters in three directions, plus an additional 4 Å.

*Vina* [34]. The software versions employed include *Vina* 1.2.3, Reduce 3.23.130521, ADFRsuite 1.0, and RDKit 2022.09.1. Ligands are processed into PDBQT files using ADFR *prepare_ligand scripts*, and RDKit calculates their centroid coordinates and coordinate ranges. Hydrogen atoms are added with Reduce, and PDBQT files are generated using the ADFR *prepare_receptor script*. The docking box size is set using the co-crystallized ligand's center and its maximum diameters in three directions, plus an additional 4 Å.

*Smina*[53]. The software version employed is *Smina* python API 2021.12.09. The ligand is inputted in SDF format, while the protein is in PDB format. Use the center of the co-crystallized ligand and *autobox_add* is set to 4 Å.

*Gnina*[54]. The software version employed is *Gnina* 1.0.3. The ligand is inputted in SDF format, while the protein is in PDB format. Use the center of the co-crystallized ligand and *autobox_add* is set to 4 Å.

DiffDock[41]. DiffDock is a docking method based on a diffusion-generated model. The protocol and settings in inference.py in the DiffDock repository ( https://github.com/gcorso/DiffDock ). The docking procedures adhere to the default settings of Diffdock.

FlexPose[42]. FlexPose is a framework for AI-based flexible modeling of protein-ligand binding pose. The protocol and settings in demo.py in the FlexPose repository ( https://github.com/tiejundong/FlexPose ) were used for docking.

For all methods, we generated 36 ligand poses and selected the one closest to the ground-truth pose for statistical analysis. To balance the computational complexity and the docking accuracy, following the previous work[51], we defined the flexible amino acids for all conventional docking algorithms as the amino acids within 3.5 Å distance from the ligand.

### S2 The Definition of Side-chain Torsion Angles.

**Table. S1.** Predefined bijective table of side-chain torsion angle with atom

| AA names | Tor_1 | Tor_2 | Tor_3 | Tor_4 |
|---|---|---|---|---|
| **Gly** | | | | |
| **Ala** | | | | |
| **Ser** | N-CA-CB-OG | | | |
| **Cys** | N-CA-CB-SG | | | |
| **Val** | N-CA-CB-CG1 | | | |
| **Thr** | N-CA-CB-OG1 | | | |
| **Pro** | N-CA-CB-CG | | | |
| **Ile** | N-CA-CB-CG1 | CA-CB-CG1-CD1 | | |
| **Leu** | N-CA-CB-CG | CA-CB-CG-CD1 | | |
| **Asp** | N-CA-CB-CG | CA-CB-CG-OD1 | | |
| **Asn** | N-CA-CB-CG | CA-CB-CG-OD1 | | |
| **His** | N-CA-CB-CG | CA-CB-CG-ND1 | | |
| **Phe** | N-CA-CB-CG | CA-CB-CG-CD1 | | |
| **Tyr** | N-CA-CB-CG | CA-CB-CG-CD1 | | |
| **Trp** | N-CA-CB-CG | CA-CB-CG-CD1 | | |
| **Glu** | N-CA-CB-CG | CA-CB-CG-CD | CB-CG-CD-OE1 | |
| **Gln** | N-CA-CB-CG | CA-CB-CG-CD | CB-CG-CD-OE1 | |
| **Met** | N-CA-CB-CG | CA-CB-CG-SD | CB-CG-SD-CE | |
| **Arg** | N-CA-CB-CG | CA-CB-CG-CD | CB-CG-CD-NE | CG-CD-NE-CZ |
| **Lys** | N-CA-CB-CG | CA-CB-CG-CD | CB-CG-CD-CE | CG-CD-CE-NZ |

### S3 Packing Dataset

**Table S2.** List of targets in Packing dataset.

| Dataset | Protein ID |
|---|---|
| **CASP** | T0950-D1,T0951-D1,T0953s1-D1,T0953s2,T0954-D1,T0955-D1,T0957s1-D1,T0957s1-D2,T0957s1,T0957s2-D1,T0958-D1,T0960,T0963,T0965-D1,T0966-D1,T0967-D1,T0968s1-D1,T0968s2-D1,T0969-D1,T0970-D1,T0971-D1,T0976,T0980s1-D1,T0984,T0990,T1003-D1,T1005-D1,T1006-D1,T1008-D1,T1009-D1,T1011,T1016-D1,T1018-D1,T1021s1-D1,T1021s2-D1,T1021s3,T1022s1,T1022s2-D1, T1024,T1025-D1,T1026-D1,T1027-D1,T1028-D1,T1029-D1,T1030,T1031-D1,T1032-D1,T1033-D1,T1034-D1,T1035-D1,T1036s1-D1,T1037-D1,T1038,T1039-D1,T1040-D1,T1041-D1,T1042-D1,T1043-D1,T1046s1-D1,T1046s2-D1,T1047s1-D1,T1047s2,T1049-D1,T1050,T1053,T1054-D1,T1056-D1,T1057-D1,T1064-D1,T1065s1-D1,T1065s2-D1,T1067-D1,T1073-D1,T1074-D1,T1079-D1,T1080- |

| | |
|---|---|
| **PDBbind** | 6qqw,6jap,6np2,6qrc,6oio,6jag,6i9a,6jb4,6seo,6jid,5ze6,6pka,6n97,6qtr,6n96,6qzh,6qqz,6k3l,6cjs,6n9l,6ott,6npp,6nsv,6n53,6eeb,6n0m,6ovz,5zcu,6mjq,6efk,6gdy,6kqi,6ueg,6qr7,6g3c,6iql,6qr4,6jib,6qto,6qrd,6e5s,5zlf,6om4,6qqv,6qtq,6os5,6s07,6mjj,6jb0,6uim,6mo0,6cjr,6uii,6sen,6kjf,6qr9,6g9f,6npi,6oip,6miv,6qts,6oi8,6c85,6qsz,6jbb,6np5,6nlj,6n94,6e13,6uil,6n92,6uhv,6q36,6qtx,6rr0,6ufo,6oiq,6qra,6m7h,6ufn,6qr0,6o5u,6ny0,6jan,6ftf,6jon,6cf7,6o9c,6qqu,6mja,6r4k,6h9v,6py0,6jaq,6k2n,6cjj,6a73,6qqt,6qre,6qtw,6np4,6n55,6kjd,6np3,6jbe,6qqq,6j9y,6h7d,6jao,6e7m,6rz6,6qtm,6miy,6jad,6mj4,6qr2,6qxa,6o9b,6ckl,6oir,6oin,6jam,6uhu,6mji,6nt2,6op9,6e4v,6a87,6cjp,6qrf,6j9w,6n93,6nd3,6os6,6dql,6qwi,6npm,6qrg,6nxz,6qr3,6qr1,6o5g,6r7d,6mo2 |

### S4 Performance of Packing experiment

**Table. S3.** Comparative evaluation of PackPocket and current methods on CASP.

| Method | Pocket Region | |
|---|---|---|
| | ANGLE ACCURACY % ↑ | ATOM RMSD Å ↓ |
| SCWRL | 66.9% | 0.632 |
| FASPR | 67.4% | 0.624 |
| AttnPacker | 76.0% | 0.391 |
| DiffPack | 81.8% | 0.369 |
| PackPocket(1) | 75.9% | 0.446 |
| PackPocket(3) | 80.6% | 0.341 |
| PackPocket(6) | 81.8% | 0.315 |
| PackPocket(10) | 85.1% | 0.271 |
| PackPocket(20) | 87.2% | 0.239 |
| PackPocket(40) | **87.8%** | **0.213** |

**Table. S4.** Comparative evaluation of PackPocket and prior methods on PDBbind.

| Method | Binding Site | |
|---|---|---|
| | ANGLE ACCURACY % ↑ | ATOM RMSD Å ↓ |
| SCWRL | 60.27% | 0.823 |
| FASPR | 57.21% | 0.876 |
| AttnPacker | 66.96% | 0.538 |
| DiffPack | 72.9% | 0.571 |
| PackPocket(1) | 71.3% | 0.565 |
| PackPocket(3) | 74.7% | 0.453 |
| PackPocket(6) | 76.1% | 0.417 |
| PackPocket(10) | 77.2% | 0.390 |
| PackPocket(20) | 78.2% | 0.361 |
| PackPocket(40) | **79.1%** | **0.336** |

### S5 Pocket Change Statistical and Experiment Number Statistical in Cross-Docking

Due to the data processing errors in different docking algorithms, the number of successful docking results varies for each method. We counted the docking test number for each method and analyzed the pocket change between the “source” and “target” proteins used for Cross-Docking tests.

**Table. S5.** Statistical analysis of the number of test sample points in cross-docking using different methods

| Method | Test number in crossdock |
|---|---|
| $PackDock$ | 7587 |
| $Vina$ | 7793 |
| $Vina_{flex}$ | 7441 |
| $Smina$ | 7818 |
| $Smina_{flex}$ | 7646 |
| $Gnina$ | 7933 |
| $Gnina_{flex}$ | 7927 |
| $Autodock$ | 7968 |
| $Autodock_{flex}$ | 7468 |
| $DiffDock$ | 7648 |
| $FlexPose$ | 6914 |

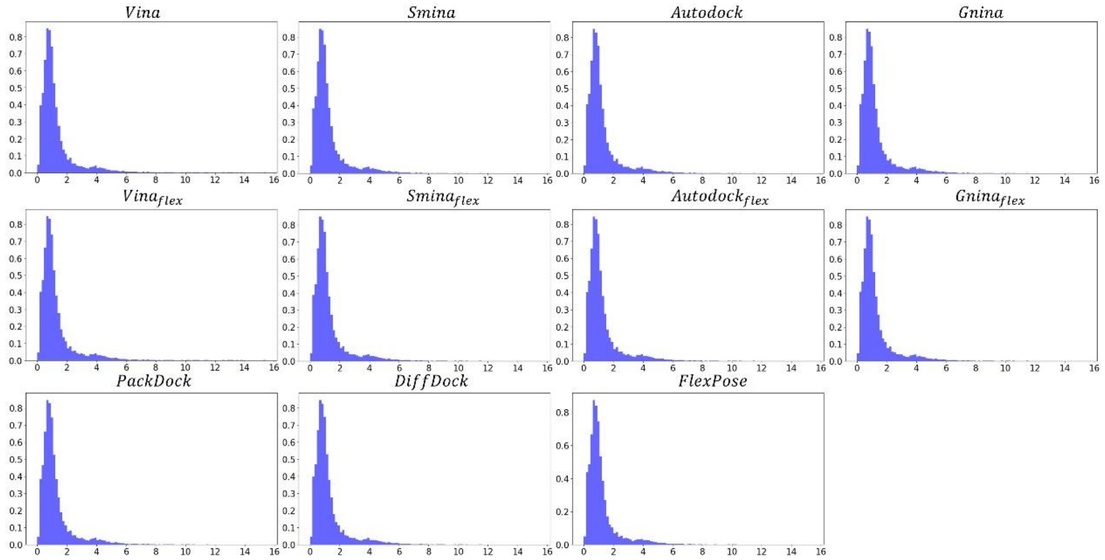


**Fig. S1.** Pocket change statistical analysis of different methods in Cross-docking testing. X-axis: Pocket change RMSD, Y-axis: Proportion

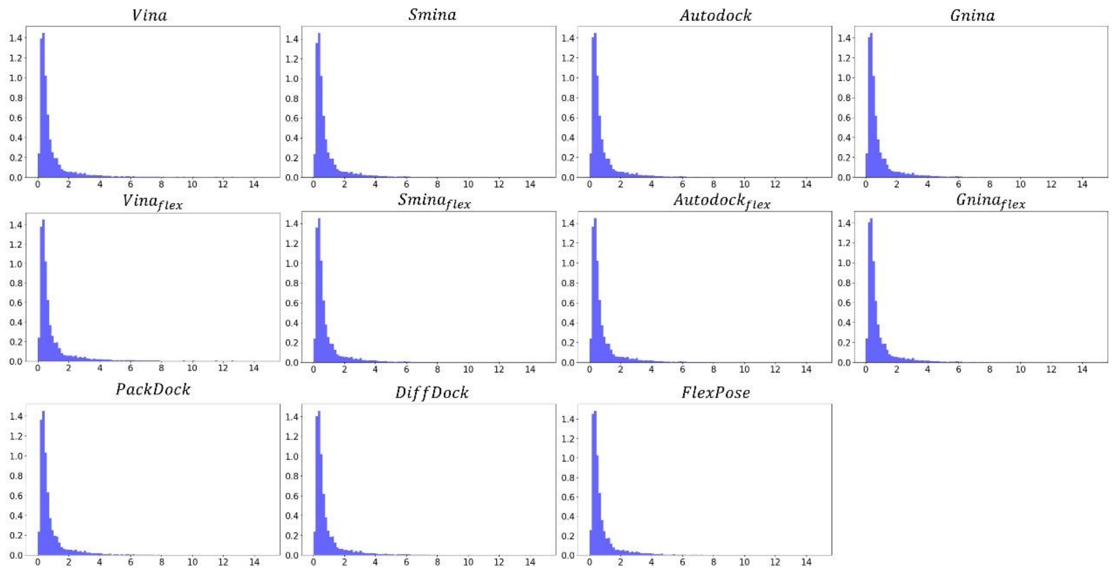


**Fig. S2.** Backbone change statistical analysis of different methods in Cross-docking testing. X-axis: Backbone change RMSD, Y-axis: Proportion

### S6 Visualization of Side-chains in PackDock

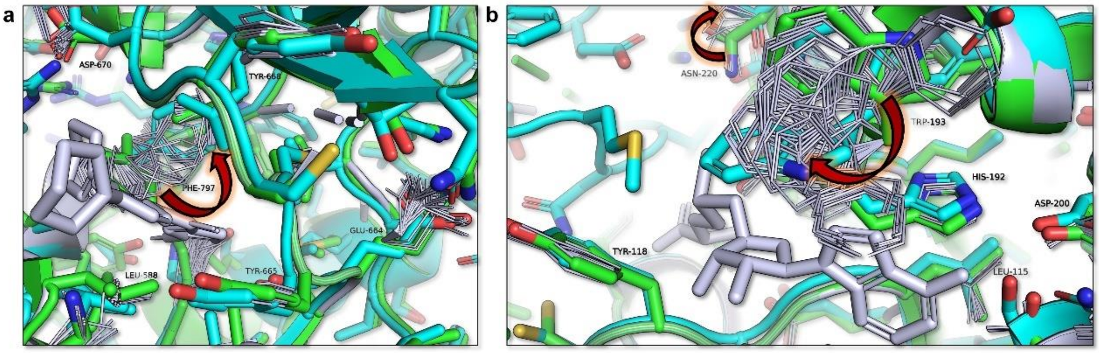


**Fig. S3.** Visualization of Side-chains in PackDock. (a) In the *apo* (2ogv, green) - *holo* (3krj, cyan) structure, F797 exhibits a significant conformational change. (b) In the *apo* (4pyi, green) - *holo* (3bmw, cyan) structure, W193 and N220 exhibits a significant conformational change. Other amino acids with smaller conformational differences show a compact unimodal distribution.
